# Close relationship between coral-associated *Chromera* strains despite major differences within the Symbiodiniaceae

**DOI:** 10.1101/825992

**Authors:** Amin R. Mohamed, Cheong Xin Chan, Mark A. Ragan, Jia Zhang, Ira Cooke, Eldon E. Ball, David J. Miller

## Abstract

Reef-building corals live in a mutualistic relationship with photosynthetic algae (family Symbiodiniaceae) that usually provide most of the energy required by the coral host. This relationship is sensitive to temperature stress; as little as a 1°C increase often leading to collapse of the association. This sensitivity has led to interest in the potential of more stress tolerant algae to supplement or substitute for the normal Symbiodiniaceae mutualists. In this respect, the apicomplexan-like microalga *Chromera* is of particular interest due to its greater temperature tolerance. We generated a *de novo* transcriptome for a *Chromera* strain isolated from a GBR coral (*“GBR Chromera”*) and compared to those of the reference strain of *Chromera* (“Sydney *Chromera*”), and to those of Symbiodiniaceae (*Fugacium, Cladocopium and Breviolum*), as well as the apicomplexan parasite, *Plasmodium falciparum*. By contrast with the Symbiodiniaceae, the two *Chromera* strains had a high level of sequence similarity evident by very low levels of divergence in orthologous genes. Although KEGG categories provide few criteria by which true coral mutualists might be identified, they do supply a molecular rationalization for the ubiquitous association of *Cladocopium* strains with Indo-Pacific reef corals. The presence of HSP20 genes may underlie the higher thermal tolerance of *Chromera*.

## Introduction

The ecological success of reef-building corals is generally attributed to their ability to establish mutualistic relationships with specific photosynthetic algae – members of the dinoflagellate family Symbiodiniaceae (formerly, the genus “*Symbiodinium*”). However, as is now widely appreciated, this relationship breaks down under environmental stress, and coral reefs globally are under threat as a consequence of the increasing frequency of weather events that exceed the thermal thresholds of corals (Hughes *et al*., 2017). In addition to Symbiodiniaceae, a wide range of uncharacterized eukaryotes are associated with corals, including apicomplexan-related lineages (ARLs) (Clerissi *et al*., 2018) that can sometimes occur in high abundance (Kwong *et al*., 2019). Two of these ARLs isolated in association with corals, *Chromera velia* (Moore *et al*., 2008) and *Vitrella brassicaformis* (Obornik *et al*., 2012), constitute the newly defined phylum Chromerida. As the closest free-living relatives of the parasitic Apicomplexa, these photoautotrophic alveolates are of considerable scientific interest (Moore *et al*., 2008). Whole genome sequencing (Woo *et al*., 2015) has recently reaffirmed the close relatedness of *Chromera* and *Vitrella*.

The nature of the relationship between corals and *Chromera* has been a subject of debate. Given its photosynthetic ability (Moore *et al*., 2008) and its ability to colonize coral larvae (Cumbo *et al*., 2013), it was initially thought that *Chromera* might be an alternative coral mutualist, potentially bringing the benefit of higher thermal tolerance than most Symbiodiniaceae (Visser *et al*., 2012; Chakravarti *et al*., 2019). However, several lines of evidence now imply otherwise. It has recently been shown that the transcriptomic response of the coral host post *Chromera* uptake (Mohamed *et al*., 2018) differed markedly from that of the same coral to a mutualistic strain of Symbiodiniaceae (Mohamed *et al*., 2016), and resembled the response to incompatible (“incompetent”) Symbiodiniaceae strains (Voolstra *et al*., 2009). The apparently hostile responses of coral larvae to *Chromera* during infection suggested that *Chromera* is more likely to be a parasite or a commensal of corals rather than a mutualist (Mohamed *et al*., 2018). Other lines of evidence support this suggestion (Barott *et al*., 2011; Janouškovec *et al*., 2012, 2013), including a recent meta-analysis which implies that *Chromera* is near exclusively associated with coral biogenous sediments (Mathur *et al*., 2018).

The present work sought to address two specific issues, in both cases making use of a *de novo* transcriptome assembly generated for a strain of *Chromera* isolated from corals on the Great Barrier Reef (GBR). Whilst *Chromera* was originally isolated from a Sydney Harbor coral, it is known to have a wide distribution (Janouškovec *et al*., 2012; Visser *et al*., 2012), and the diversity within this monospecific genus has not been systematically explored. Given the metabolic diversity that is now known to exist (LaJeunesse *et al*., 2018) within what was previously known as “*Symbiodinium*”, the extent to which conclusions about the coral-*Chromera* interaction based on the GBR isolate are generalizable is unknown.

The first goal of the present study was therefore to estimate the degree of divergence between the GBR strain of *Chromera* and that originally isolated from Sydney harbor. Given the evidence that *Chromera* is unlikely to be a coral mutualist, the second goal was to investigate the repertoires of genes that are thought to play roles in symbiosis and environmental stress tolerance in *Chromera* and compare these with those of three members of the Symbiodiniaceae. *Cladocopium goreaui* (formerly Clade C1 *Symbiodinium*) was isolated from a colony of *Acropora tenuis* on the GBR (Howells *et al*., 2012), and is mutualistic with many Indo-Pacific corals, particularly *Acropora* species. *Breviolum minutum* (formerly Clade B *Symbiodinium*) was isolated from the Caribbean coral *Orbicella faveolata*, and is a mutualist of Caribbean corals. *Fugacium kawagutii* (formerly Clade F *Symbiodinium*) was originally isolated in association with the Hawaiian reef-building coral *Montipora verrucosa* and the initial whole-genome analyses followed the assumption that *Fugacium* is a coral mutualist (Lin et al. 2015). However, *Fugacium* failed to infect juvenile corals (Yuyama *et al*., 2016), and the consensus now is that *Fugacium* is probably a surface associate of corals rather than an endosymbiont (Liu *el al*., 2018; LaJeunesse *et al*., 2018; González-Pech *et al*., 2019). Thus, the expectation was that, with respect to metabolic repertoire, the *Chromera* strains would resemble *Fugacium* rather than the known coral mutualists, *Cladocopium* and *Breviolum*. Whilst the results suggest that small HSPs may account for the tolerance of *Chromera* to elevated temperatures, they were inconclusive with respect to the nature of relationships between corals and *Chromera* or *Fugacium*. The comparative analyses do, however, provide a molecular rationalization for the near ubiquitous association of *Cladocopium* with Indo-Pacific corals in general and with *Acropora* spp in particular.

## Experimental procedures

### Chromera culture and culturing conditions

A culture of *Chromera* (Mdig3 strain) from the University of Technology Sydney (Cumbo *et al*., 2013) was used in this study and referred to as “GBR *Chromera”*. The identity of the culture was confirmed both by microscopy and by using *Chromera*-specific PCR primers (Supplementary information Figure 1, Table 1). This *Chromera* strain was originally isolated from the stony coral *Montipora digitata* (Acroporidae) from Nelly Bay, Magnetic Island on the inner central part of the Great Barrier Reef. Cultures were maintained at 25 °C in Guillard’s f/2 medium on a 12 h/12 h day and night regime. Note that *Chromera* was subjected to a variety of treatments prior to RNA extraction in order to ensure that the transcriptome assembly captured as many genes as possible. Culture conditions included control, dark stress, cold shock, heat shock, motile and mixotrophic (for details see Supplementary Methods). In all cases, exponentially growing cultures were separated and subjected to the treatment condition and harvested at the end of the experimental treatment. During culturing no antibiotics were used to exclude any potential contribution of the antibiotic treatment to the mRNA expression in the cultures.

**Table 1.** Illumina sequencing and mapping statistics: number of reads and bases of raw and processed data after quality control (adapter removal, trimming and filtering low-quality bases) for each library and percentages of reads successfully mapped onto the GBR *Chromera* transcriptome. Note that *Chromera* was subjected to a variety of treatments prior to RNA extraction in order to ensure that the transcriptome assembly captured as many genes as possible.

|  | RNA-Seq library/ culturing condition |  |  |  |  |  |  |  |
| --- | --- | --- | --- | --- | --- | --- | --- | --- |
|  | Control | Cold | Heat | Dark | Motile | Mixotrophic | Total | Average |
| No. raw paired-end reads (M) | 31.35 | 31.75 | 31.49 | 31.8 | 30.17 | 32.91 | 189.47 | 31.57 |
| No. raw bases (Gb) | 6.27 | 6.35 | 6.29 | 6.36 | 6.03 | 6.58 | 37.88 | 6.31 |
| No. reads passed QC (M) | 27.59 | 28.03 | 27.46 | 27.9 | 26.47 | 28.93 | 166.38 | 27.73 |
| No. bases passed QC (Gb) | 5.51 | 5.6 | 5.49 | 5.58 | 5.29 | 5.78 | 33.25 | 5.54 |
| Mapping rate (%) | 85.2 | 83.9 | 86.9 | 83.7 | 81.3 | 82.5 | - | 84.2 |

M= million; Gb= gigabase

### RNA isolation and high-throughput sequencing

50 mL of *Chromera* cultures were pelleted by spinning the cultures at 3,000 x *g* for 5 min. Pellets were suspended in 1 ml 0.2 μm sterile FSW and centrifuged at 3,000 x *g* for 5 min. Pellets were snap frozen in liquid nitrogen and stored at −80°C until further treatment. Total RNA was isolated from ∼80 mg of the frozen *Chromera* pellets using the RNAqueous® Total RNA Isolation Kit (Ambion). The pellets were lysed twice for 20s at 4.0 ms^-1^ in Lysing Matrix D tubes (MP Biomedicals, Australia) containing 960 µL of lysis/binding solution plus 80 µL of the Plant RNA Isolation Aid (Ambion, USA) on a FastPrep®-24 Instrument (MP Biomedicals, Australia). RNA was bound to filter cartridges supplied with the kit and washed three times, finally RNA was eluted in 40 µL of the elution solution. RNA quantity and quality were assessed using a NanoDrop ND-1000 spectrometer, Qubit® 2.0 fluorometer and Agilent 2100 bioanalyzer. Messenger RNA (mRNA) was isolated from 1 µg of total RNA and 6 RNA-Seq libraries were prepared using the TruSeq RNA Sample Preparation Kit (Illumina). Libraries were sequenced on an Illumina HiSeq 2000 platform at the Australian Genome Research Facility (AGRF) in Melbourne, Australia. Sequencing produced a total of 189.5 million individual 100 bp paired-end reads (Table 1).

### Processing of Illumina data

The raw Illumina reads were filtered and adapters were clipped using TRIMMOMATIC (v0.32) (Bolger *et al*., 2014). Reads were filtered based on quality and size as follows; both universal and indexed Illumina adapters were clipped, quality trimming was also performed by removing leading and trailing bases with Phred quality score < 25 and average Phred quality score was calculated in 4 bp sliding windows. Bases were trimmed from the point in the read where average Phred quality score dropped below 20 (i.e. the chances that a base is called incorrectly is 1 in 100) and reads of < 50 bp were also excluded.

### De novo assembly and annotation of transcriptome

The trimmed/filtered Illumina reads were used for *de novo* transcriptome assembly using Trinity (r20140717 version). The assembly was carried out with the recommended protocol described in (Haas *et al*., 2013) and using options appropriate for *de novo* transcriptome assembly of strand specific RNA-Seq libraries. Minimum contig length of 500 and read normalization were specified. Trinity collects transcripts with shared sequence identity into clusters that are loosely related to genes. The longest isoform per cluster was selected using a custom Perl script from the assembled Trinity output “assembled transcriptome” for the purpose of annotation. *Chromera* contigs were annotated by similarity search using batch BLASTX conducted locally against the Swiss-Prot protein database downloaded in September 2014 (E-value cut off 10^−3^ and maximum 20 hits). Raw BLASTX outputs were imported to Blast2GO suite (version 2.6.5) (http://blast2go.com/b2ghome) for functional annotation and Gene Ontology (GO) assignment. KEGG analysis was also performed using the KEGG Automatic Annotation Server (KAAS) (Moriya *et al*., 2007) (http://www.genome.jp/kaas-bin/kaas_main) in order to obtain an overview of the associated metabolic pathways. The bi-directional best hit (BBH) method was used to obtain KEGG orthology (KO) assignments.

In order to validate the accuracy of the *de novo* assembly, reads were mapped back to the *de novo*-assembled transcriptome using the BOWTIE aligner version 0.12.7 (Langmead and Salzberg, 2012) with default mapping parameters. The percent of the mapped reads as proper pairs was used to assess the assembly quality. Moreover, BLASTN (E-value of ≤ 10^−10^) was performed against bacterial genomes downloaded from the GenBank, NCBI to determine the percentages of putative bacterial transcripts in the dataset. The completeness of the transcriptome assembly was assessed using BUSCO v3.1.0 making use of the function *run_BUSCO*.*py* (--mode transcriptome) with the Eukaryota_odb9, alveolate_stramenophile and protists_ensembl data (retrieved 23 October 2019).

### Phylogenomic analyses

A multi-gene phylogenetic analysis was performed to assess the relative phylogenetic distance between *Chromera* strains and infer their evolutionary relationship to other Apicomplexans and Symbiodiniaceae. The protein sequences of GBR *Chromera* were predicted by Transdecoder (Haas *et al*., 2013). Transcript nucleotide (CDS) and protein sequences for *Chromera* CCMP2878 strain “Sydney *Chromera*” were downloaded from CryptoDB (release-37; http://cryptodb.org/cryptodb/), and for *Plasmodium* falciparum from PlasmoDB (release-37; http://plasmodb.org/plasmo/). Corresponding data for Symbiodiniaceae species were based on gene models from their respective genome sequencing projects; specifically, *Fugacium kawagutii* (Lin et al 2015) was obtained from the Symka Genome Database (http://web.malab.cn/symka_new/download.jsp), *Cladocopium goreaui* (Liu et al 2018) from ReFuGe 2020 site (http://refuge2020.reefgenomics.org/) and *Breviolum minutum* (Shoguchi *et al*., 2013) from the OIST Marine Genomics online resource (https://marinegenomics.oist.jp/symb/viewer/info?project_id=21).

The longest transcript for each gene was extracted for all six species and used to infer orthologous clusters with OrthoFinder (Emms and Kelly, 2015). A total of 692 orthogroups were found to have representative genes in all species, and of these 172 consist of single-copy genes in each species; they represent strictly orthologous gene sets. Amino acid sequences for these 172 orthologous sets were aligned using MAFFT (Katoh and Standley, 2013) and converted into corresponding codon alignment by pal2nal (Suyama *et al*., 2006). Poorly aligned regions were removed using trimAl based on distribution of gaps (Capella-Gutierrez *et al*., 2009). Finally, IQ-TREE (Nguyen *et al*., 2015) was used to perform a partitioned phylogenetic analysis allowing independent estimation of evolutionary model for each protein set. To summarise the effect of phylogenetic relatedness on overlap between gene repertoires the number of shared orthologs was calculated for all members of each clade based on orthogroup information generated with OrthoFinder (see above). Additionally, for each pair of taxa the orthogroup information was used to compute a Hammings distance based on the ratio of the number of unshared orthogroups (those unique to either taxon in the pair) to the total number of orthogroups shared by the pair.

### Comparative transcriptomics

Transcriptomes of Sydney *Chromera, Fugacium*, and the known mutualistic Symbiodiniaceae (*Cladocopium* and *Breviolum)* were also mapped against the KEGG database as previously performed for the GBR strain (using the bi-directional best hit (BBH) method). Genes mapped to different KEGG categories were calculated and compared. To better understand relationships between corals and both *Chromera* and *Fugacium*, repertoires of genes in categories that are important for symbiosis, such as ABC transporters, as well as those involved in processes such as nitrogen metabolism and stress tolerance, were compared. The overlap amongst these genes was plotted using the R package UpSetR https://github.com/hms-dbmi/UpSetR/ (Conway *et al*., 2017).

## Results and discussion

### GBR Chromera transcriptome assembly and annotation

After confirming the identity of *Chromera* cultures using novel *Chromera*-specific PCR primers, a transcriptome assembly was generated from 166 million paired-end Illumina reads (∼33 million per library; Table 2). The number of putative genes (39 457, based on the longest transcript isoform per Trinity gene cluster) identified in the GBR *Chromera* isolate (Table 2) is comparable to those predicted for various Symbiodiniaceae isolates (30 000-49 000) based on transcriptome and genome data (Bayer *et al*., 2012;Shoguchi *et al*., 2013;Rosic et al., 2015;Aranda *et al*., 2016; Liu *et al*. 2018 and Shoguchi *et al*., 2018), but is higher than the number predicted from the *Chromera* genome of the reference strain (26 112 excluding TEs; Woo *et al*., 2015). As only 19.4% of the GBR *Chromera* genes had significant BLASTX hits against the Swiss-Prot protein database, the majority of *Chromera* genes code for unknown functions. This level of novelty is to be expected for organisms such as chromerids that are evolutionarily distant from well-characterized species, and has previously also been observed with Symbiodiniaceae and other dinoflagellates (Lin *et al*., 2010;Bayer *et al*., 2012; Stephens *et al*., 2018). To assess the quality of the *de novo* assembled transcriptome, reads were mapped to the assembly and an average of 84% of the paired Illumina reads were mapped successfully (Table 1). The assembled transcriptome was judged to be relatively comprehensive on the basis of high percentages of reads mapping.

**Table 2.** Summary statistics for the GBR *Chromera de novo* transcriptome assembly using Trinity and annotation based on Swiss-Prot (SP), Gene Ontology (GO) and KEGG databases

|  |  |
| --- | --- |
| **Trinity outputs:** |  |
| Total Trinity 'genes' | 39 457 |
| Total Trinity transcripts | 79 842 |
| GC % | 53.42 |
| **Statistics based on ALL transcript contigs:** |  |
| Contig N50 | 2 289 |
| Median contig length | 1 461 |
| Average contig | 1 838.03 |
| Total assembled bases | 146 752 195 |
| **Statistics based on ONLY LONGEST ISOFORM per 'GENE':** |  |
| Contig N50 | 2 220 |
| Median contig length | 1 403 |
| Average contig | 1 764.26 |
| Total assembled bases | 69 612 282 |
| **Annotation statistics based on longest isoform per gene data:** |  |
| Swiss-Prot (SP) database | 7 644 |
| Gene Ontology (GO) database | 5 225 |
| KEGG database | 4 220 |

5225 of Swiss-Prot annotated genes (68.35%) were assigned to 38271 GO terms. Biological process (GO-BP) accounted for the majority of GO terms (22 205, 58.02%), followed by cellular component (GO-CC; 11 149, 29.1%) and molecular function (GO-MF; 4 917, 12.8%). Functions involved in *cellular process* and *metabolic process* (16% and 14%, respectively) were highly represented amongst GO-BP. In GO-MF, the most represented terms were *catalytic activity* (46%) followed by *binding* (36%). In GO-CC, the terms *cell* (39%) and *organelle* (33%) were highly represented (Supporting information Figure 2). Most of the KEGG-based annotations (34% of all assignments) were assigned to the metabolic pathway category, followed by the human disease category (18% of all assignments) (Supporting information Figure 3). Moreover, signal transduction and infectious diseases were the most highly represented pathways (Supporting information Figure 4, Table 2).

Several different approaches were used in order to assess the completeness of the assembled transcriptome. The KEGG annotation was searched for essential protein complexes/ pathways and the majority of genes for the pathways were found. Searched complexes included core cellular/ molecular protein complexes and pathways such as Ribosome biogenesis in eukaryotes, Ribosome, RNA polymerase, Spliceosome and Proteasome (Supporting information Table 3 and Figures 5-9). The use of BUSCO analysis resulted in a moderately high recovery (∼60%) of conserved eukaryotic genes and relatively low recovery (∼26-46 %) of conserved alveolate and protist genes, respectively, using default settings (Supporting information Table 4). In addition, 0.1% of the assembled *Chromera* sequences (contigs) had BLASTN hits to bacterial databases (E-value ≤ 10^−10^) indicating very low bacterial contamination.

### Sequence and functional similarities of the two Chromera strains

Phylogenomic analyses based on 172 orthologous single copy genes revealed a close relationship (branch length 0.07) between the GBR and Sydney strains of *Chromera* compared with divergences between different species within the Symbiodiniaceae (branch lengths 1.23 to 2.27) (Fig.1). Pairwise nucleotide similarity between the Sydney and GBR *Chromera* isolates was 96.2%, whereas the corresponding figures for the genera of Symbiodiniaceae were 72-78%. A close relationship between the *Chromera* isolates was further supported by the presence of a relatively large number (14645) of shared orthologous genes and an average nucleotide identity of 99.12% based on alignments between these one-to-one orthologs. The overall distributions of the six main KEGG categories were similar in the two *Chromera* strains, one third of KEGG-annotated genes being assigned to *metabolism* (Supporting information Figure 10). This might reflect broadly similar functions and lifestyles. The only attempt to understand the nature of the coral-*Chromera* association used the Sydney strain (Mohamed *et al*., 2018). Given the high level of similarity between the GBR and Sydney harbour strains, the responses of corals to them are unlikely to differ significantly.

**Figure 1.**
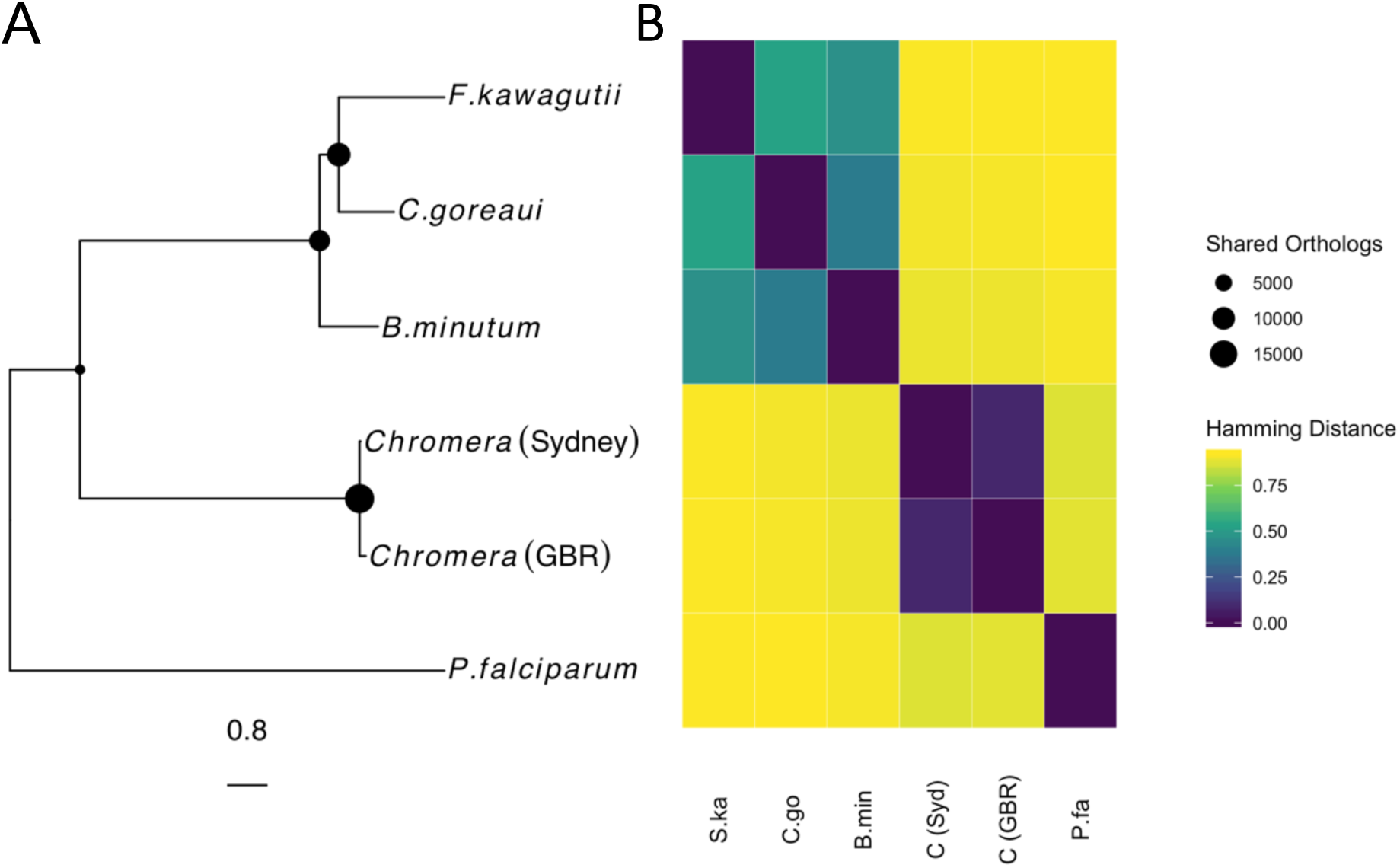
(A) Unrooted phylogenetic tree based on maximum-likelihood phylogenetic analysis of 172 orthologous nucleotide sequences from Sydney *Chromera*, GBR *Chromera, P. falciparum, F. kawagutii, B. minutum* and C. *goreaui*. All nodes in the tree were fully supported (100% of bootstraps) based on 1000 bootstraps by ultrafast bootstrap in IQtree. The sizes of filled circles at some nodes in the tree indicate numbers of shared othologs. (B) Heat map summarizing Hamming distances between taxa based on nucleotide similarity in the 172 orthologous sequences used for phylogeny reconstruction.

### Focus on genes implicated in the symbiotic lifestyle

To gain additional perspectives on the coral-*Chromera* interaction, the assembled transcriptomes for the *Chromera* strains were compared to those for *Fugacium* (which is assumed to be non-mutualistic) and to those of *Cladocopium* and *Breviolum*, which are mutualistic Symbiodiniacae strains, focusing on categories of genes likely involved in symbiosis, particularly nitrogen metabolism, transport and stress.

### Metabolic exchanges in coral-algal symbioses

Nutrient exchange between the symbiotic partners is of major importance in coral-*Symbiodinium* mutualisms (Meyer and Weis 2012, Davy *et al*., 2012, Lin *et al*., 2015, Aranda *et al*., 2016). However, the nature of the translocated material(s) and mechanisms underlying exchanges between the coral host and algal symbionts are still unclear. To assess the potential for nutrient exchange between *Chromera* and coral hosts, the representation of the KEGG pathways *Nitrogen metabolism* and *ATP-binding cassette (ABC) transporters* were investigated in the two *Chromera* strains and compared to those in the three members of the Symbiodiniaceae (*Fugacium, Cladocopium and Breviolum*).

The *Nitrogen metabolism* category was investigated on the basis that nitrogen cycling or conservation appears to be critical to the coral-*Symbiodinium* mutualism. Nitrogen is thought to be a growth limiting factor in nutrient-poor tropical waters, and many marine microbes have the ability to assimilate inorganic nitrogen (Pernice *et al*., 2012), for a recent review see Radecker *at al*. (2015). The results of a survey of genes captured under the KEGG pathway identifier KO00910 (Nitrogen metabolism) are presented in Supporting information Table 4. Note that this KEGG pathway does not include ammonium transporters, although it has previously been reported that the genomes of Symbiodiniaceae encode multiple ammonium transport proteins (Aranda *et al*., 2016).

In ocean waters, nitrate is generally the most abundant form of available nitrogen, concentrations often being at least an order of magnitude higher than those of ammonium and nitrite. However, *in hospite*, ammonium is likely to be the dominant nitrogen source available to intracellular symbionts. Thus, facultative symbionts (as most Symbiodiniaceae are thought to be) must not only be nutritionally versatile, but also able to regulate genes involved in the transport and assimilation of different N-sources. As might be expected, all of the algae surveyed encode nitrate /nitrite transporters as well as enzymes required for assimilation of nitrogen in these forms. Although nitrate/nitrite transporters (NRTs) were present in all of the algae surveyed, some differences were apparent. Whereas the NRTs of *Breviolum, Fugacium* and *Chromera* were the MFS-type (K02575), this type was not detected in *Cladocopium*; rather, in this organism, components (K15577 and K15579) of a distinct ABC-type NRT were found. This difference may be significant; in cyanobacteria, the MFS-type has high affinity for both nitrate and nitrite, whereas the ABC-type has a much higher affinity for nitrate (Maeda and Omata, 2009).

### ATP-binding cassette (ABC) transporters

The ABC class includes the largest number of transporters involved in either or both uptake and export of a wide range of substrates, including inorganic ions, carbohydrates and lipids. ABC transporters have been implicated in translocation of nutrients and metabolites in cnidarian symbioses (Davy *et al*., 2012; Mathews *et al*., 2017; Mohamed *et al*., 2019), and hence were an obvious focus for comparative analyses.

As with the nitrogen metabolism category, surveying ABC transporter complements (Fig. 2) was largely unsuccessful in providing general molecular criteria by which known mutualists (*Breviolum* and *Cladocopium*) can be distinguished from commensals or parasites. One potentially significant difference, however, is that both *Breviolum* and *Cladocopium* encode K10111 members, which are nominal transporters of a variety of different sugars and which may be involved in carbohydrate translocation *in hospite*, whereas proteins of this type were not detected in either *Fugacium* or *Chromera* (Supporting information Table 5). Unfortunately, the survey provides few other grounds for speculation about photosynthate translocation. Whilst a number of candidates for roles in sterol or lipid translocation were detected, these were generally not restricted to the coral mutualists. For example, components of the mla/lin type transport system for phospholipids/sterols/gamma-HCH (K02065 and K02066) were detected in *Fugacium* as well as *Cladocopium*, and the *Chromera* ABC repertoire does include possible sterol transporters (K05683, K05681, K08712).

**Figure 2.**
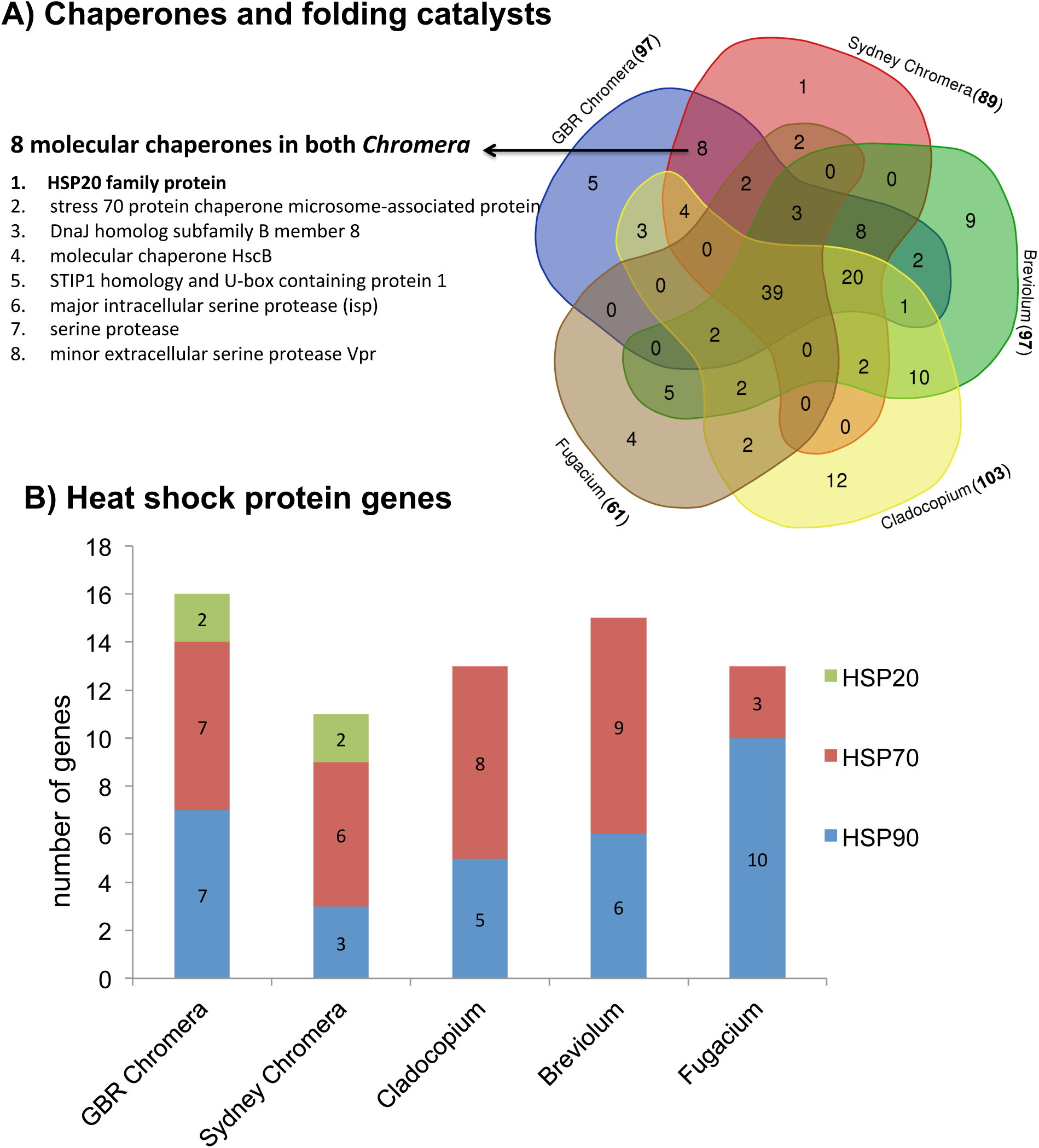
UpsetR plot (which allows comparative visualisation of the presence/absence of genes of a pathway in different taxa) illustrating the intersection between genes of two KEGG pathways: (A) Nitrogen metabolism (KO00910) and (B) ABC transporters (KO02010) in the two *Chromera* strains, *Fugacium*, mutualistic Symbiodinianceae (*Cladocopium* and *Breviolum*) and the parasite *P. falciparum*. Each plot is generated from a binary matrix with unique KEGG identifiers for each pathway and each taxon in a column. The number of genes per taxon is represented by the length of the orange bars. The black dots represent presence/absence. The purple columns show the number of genes shared amongst taxa or exclusive to one of them. So, reading the upper plot from the left there are 8 different genes (each gene is represented once) that are exclusively found in *Cladocopium* and not the others, second column shows 3 genes that are found in the 3 symbiodiniaceae and the 2 *Chromera*, but not present in *P. falciparum*, the third column shows 3 genes that are found in all taxa considered, etc. The intersection size reflects the number of unique or shared genes.

Despite these analyses being uninformative on the issue of general characteristics of mutualists, they do provide grounds for speculation about the near-ubiquitous association of *Cladocopium* with Indo-Pacific corals (LaJeunesse *et al*., 2018) and particularly with *Acropora* spp.. The ABC repertoire of *Cladocopium* by far exceeded those of all of the other organisms surveyed (Fig. 2), and included many genes identified only in this organism, amongst which were transporters for sugars, lipids and amino acids as well as inorganic nutrients. A range of candidate amino acid transporters were restricted to *Cladocopium*, one of which (HisP) is of particular interest in the context of coral symbioses, as it encodes a histidine transport ATP-binding protein (Supporting information Table 6). The *Cladocopium* strain included here was originally isolated from *Acropora tenuis* (Liu *et al*., 2018), a member of the complex coral superfamily, and whereas robust corals (members of the other coral superfamily) have a complete histidine biosynthetic pathway, complex corals (like all other animals) must acquire this either from the diet or from their resident endosymbionts (Ying *et al*., 2018). Also intriguing in the context of mutualism is the detection only in *Cladocopium* of three components required for cystine transport (K02424, K10009, K10010); the interest in these stems from the fact that, amongst corals, members of the genus *Acropora* lack one of the enzymes required for cysteine biosynthesis (Shinzato *et al*., 2011). In this respect, *Acropora* appears to be unique; all other corals surveyed had a compete cysteine pathway. The presence of an efficient cystine transport system may therefore underlie the near ubiquitous association of *Cladocopium* C1 with *Acropora* spp.

### Fugacium – an evolutionary recidivist?

The ABC transporters identified in *Fugacium* include several candidates for roles in nutrient exchange – for example, K05641 encodes an ABCA1 protein known as the cholesterol efflux regulatory protein (CERP) that mediates efflux of cellular cholesterol and phospholipids (reviewed by Zhao and Lappalainen, 2012). Note that an ABCA1 gene was up-regulated in adult *Acropora millepora* colonies in the light, i.e. when transport of photosynthate from alga to host occurs (Bertucci *et al*.,2015). K02065, which was present in all three Symbiodiniaceae, encodes a phospholipid/cholesterol/gamma-HCH transport system ATP-binding protein, again with a potential function in lipid/sterol translocation. Indeed, some ABC family A transporters (ABCA7 and ABCA3) were upregulated in *Cladocopium* during establishment of symbiosis with coral larvae (Mohamed *et al*., 2019). Thus, although *Fugacium* is now considered not to be a coral mutualist, the comparative analyses presented here do not rule this out. Whilst the nature of the relationship between *Fugacium* and corals is still unclear, the apparent contradictions could, however, be rationalized by consideration of the phylogenetic position of *Fugacium*. The ability to become endosymbiotic is considered to be a defining characteristic of the family Symbiodiniaceae (LaJeunesse *et al*., 2018); as a member of a derived clade within the Symbiodiniaceae, *Fugacium* may have undergone (or be undergoing) secondary loss of symbiotic potential but its genome still bear vestiges of this ancestral condition.

### The small heat shock protein HSP20 might explain stress tolerance in Chromera

Molecular chaperones are important for refolding damaged proteins (Vierling, 1991), thus their involvement in stress tolerance is inevitable. Amongst these chaperones we looked at heat shock proteins in both *Chromera* and two symbiotic members of Symbiodiniaceae, *Cladocopium* and *Breviolum*. Despite genes coding for chaperones in the KEGG Orthology 03110 *Chaperones and folding catalysts* (including HSP90/70) being present in similar numbers in all of these genera, gene(s) coding for two HSP20 isoforms were found in both *Chromera* strains to the exclusion of the three Symbiodiniaceae algae (Fig. 3). Consistent with a potential role in stress tolerance the HSP20 was reported recently to be correlated with stress tolerance in reef-building corals where coral species containing more HSP20 genes were more stress tolerant (robust corals) than those with smaller numbers of HSP20 genes (complex corals) (Ying *et al*., 2018).

**Figure 3.**
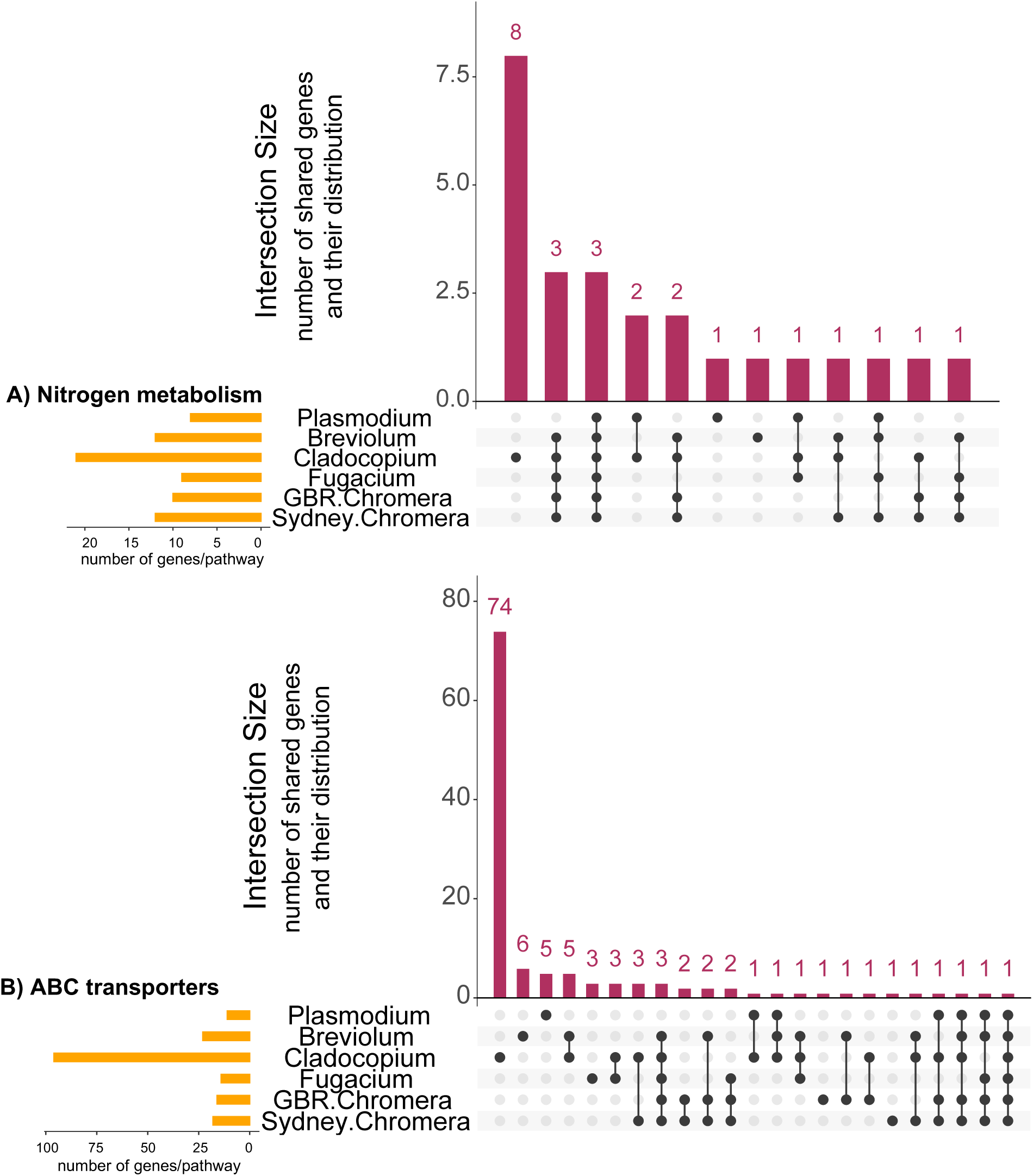
Comparison of the repertoires of genes encoding chaperones and folding catalysts (KO03110) of *Chromera* strains and Symbiodiniaceae. (A) A five-way Venn diagram of the gene overlap amongst the two *Chromera* strains, mutualistic Symbiodiniaceae (*Cladocopium* and *Breviolum*) and *Fugacium*. (B) A bar graph illustrating numbers of HSP20, HSP70 and HSP90 genes identified in the five algal datasets.

## Conclusions

This paper describes *de novo* assembly and annotation of a transcriptome for a *Chromera* strain isolated from a GBR coral, which was generated in order to understand diversity within the species and to provide a better understanding of the metabolic capabilities and lifestyle of this photosynthetic apicomplexan alga. The GBR strain has a high degree of similarity with the Sydney (culture collection) strain, hence infection studies based on the former (e.g. Mohamed *et al*., 2018) can be generalized to the latter. Comparisons between *Chromera*, the Symbiodiniaceae algae *Fugacium, Cladocopium* and *Breviolum* and the apicomplexan parasite *P. falciparum* based on specific categories of genes were inconclusive with respect to common molecular characteristics of mutualists, but suggest that HSP20 genes may underlie the higher thermal tolerance of *Chromera* compared to Symbiodiniaceae. The presence of specific genes implicated in mutualism suggest that *Fugacium* may have secondarily lost the ability to establish symbioses, and the comparative analyses provide a molecular rationale for the near-ubiquitous association of *Cladocopium* with Indo-Pacific corals.

## Acknowledgments

The research was supported by the Australian Research Council through Grant CE140100020 to DM via the ARC Centre of Excellence for Coral Reef Studies at James Cook University. AM was supported by PhD scholarships provided by the Cultural Affairs and Mission sectors of the Egyptian Ministry of Higher Education and Scientific Research and James Cook University Postgraduate Research Scholarship (JCUPRS) and through the AIMS@JCU joint-scheme. The authors gratefully thank Dr Vivian Cumbo at Macquarie University for supplying *Chromera* cultures.

## DATA AVAILABILITY

Data reported in this study have been submitted to the NCBI Gene Expression Omnibus (GEO) under accession number GSE139820 including raw illumina data, transcriptome assembly and annotations.

## Author contributions

AM and DM conceived and designed the study. AM conducted the experiments and generated the RNA-Seq data. AM performed bioinformatics required for transcriptome assembly, annotation and comparative analyses with guidance from CXC and MR. JZ and IC performed the phylogenomic analysis. AM, EB and DM interpreted that data and wrote the manuscript. All authors read the article and approved the final version.

