## Supporting info for "Close relationship between coral-associated *Chromera* strains despite major differences within the Symbiodiniaceae"

**Supplementary Methods**

**Identity check of *Chromera* cultures**

***Microscopic check of Chromera culture***

The starting *Chromera* culture was checked with an inverted microscope for protist and bacterial contamination before small aliquots were subjected to genetic identification, growth and further application of the experimental treatments.

*Genetic Identification*

***Chromera gDNA extraction***

gDNA was extracted from 50 ml of exponentially growing culture. Cultures were centrifuged at 9000 rpm for 5 minutes at 4 °C, the Chromera pellet was resuspended in 1ml fresh f/2 medium, centrifuged at maximum speed for 5 minutes at 4°C and stored at -80 °C until further treatment. The ISOLATE II Plant DNA Kit (BIOLINE) was used for DNA extraction according to the manufacturer’s instructions. DNA was eluted in 100 μl of elution buffer in a 1.5 ml tube. DNA was checked by running onto an agarose gel and a Nanodrop ® ND-100 Spectrophotometer (Wilmington, U.S.A) was used to estimate the concentration and quality of the DNA obtained from the DNA extractions. Milli-Q water was used to blank the instrument. 1.5 μl of sample was placed directly onto a fibre optic measurement surface where a retention system using surface tension held the sample in place. DNA concentrations, absorbance at 230 (λ230) and the ratio 260/280 were recorded.

***Polymerase chain reaction (PCR) Primer Design***

*Chromera* large subunit ribosomal RNA gene, partial sequence, GenBank: EU106870.1, (Moore *et al.*, 2008) and *C. velia* clone JS497 18S ribosomal RNA gene, partial sequence; internal transcribed spacer 1, 5.8S ribosomal RNA gene, and internal transcribed spacer 2, complete sequence; and 28S ribosomal RNA gene, partial sequence, GenBank: JN935835.1 (Morin-Adeline *et al.*, 2012) were retrieved from GenBank and used as templates for designing the PCR primers. The following primers (Table 1) were used to check the identity of the starting culture.

***Amplification of Chromera ribosomal genes using PCR***

Amplification of *Chromera* ribosomal genes was undertaken using specific primers (above) to obtain a PCR product ranging between 416 to 778 bp in size. PCR reaction was conducted in 50 μl using 1 μl of *Chromera* gDNA (approx. 100ng of DNA) as template. 1 μl of GoTaq® DNA polymerase and 2X GoTaq® reaction buffer and, 5 μl of each primer were used and finally sterile MQW was added to the reaction mixture to make a total volume of 50 μl. The PCR profile was one cycle for 2 min at 94 °C for initial denaturation followed by 34 cycles of 30 sec at 94 °C, annealing for 30 sec at 47 °C/ 51 °C and extension for 2 min at 72 °C. The final extension was at 72 °C for 10 min. The obtained amplicons were run on 1.5% Agarose gel and visualized using a UV trans-illuminator.

***Chromera culture and culturing conditions***

Cultures growing in the mid exponential (log) phase (+11 days after inoculation) were harvested at the middle of the cultures’ daytime phase and labeled as “control”. In order to maximize the variety of expressed genes, the cultures were subjected to a variety of treatments before RNA isolation and preparation of cDNA libraries. Cultures were subjected to dark stress (24 hour dark period), cold shock (4°C for 4 hours) and heat shock (36°C for 4 hours). Cultures growing in the control conditions +8 days after inoculation cultures were harvested at the middle of the cultures’ daytime phase and labeled as “motile” as cultures showed both *Chromera* life forms. In addition, cultures were also grown in f/2 media autotrophically while supplemented with exogenous organic compounds at final concentration of total 0.1%(w/v) including; Galactose (D+)  (D00201; Sigma-Aldrich), sodium acetate (D00385; ICN Biomedicals) and Glycerol (D00217; Sigma-Aldrich) and labeled as “mixotrophic”. In all cases, exponentially growing cultures were separated and subjected to the treatment condition and harvested at the end of the experimental treatment. During culturing no antibiotics were used to exclude any potential contribution of the antibiotic treatment to the mRNA expression in the cultures.

***Illumina data quality check***

Illumina raw reads from each paired end file were visualized using FASTQC version 0.11.2 in order to determine the quality of the data. In addition, reads were inspected for adapter contamination by searching for the IIlumina universal and indexed adapters.

**Supplementary Figures**

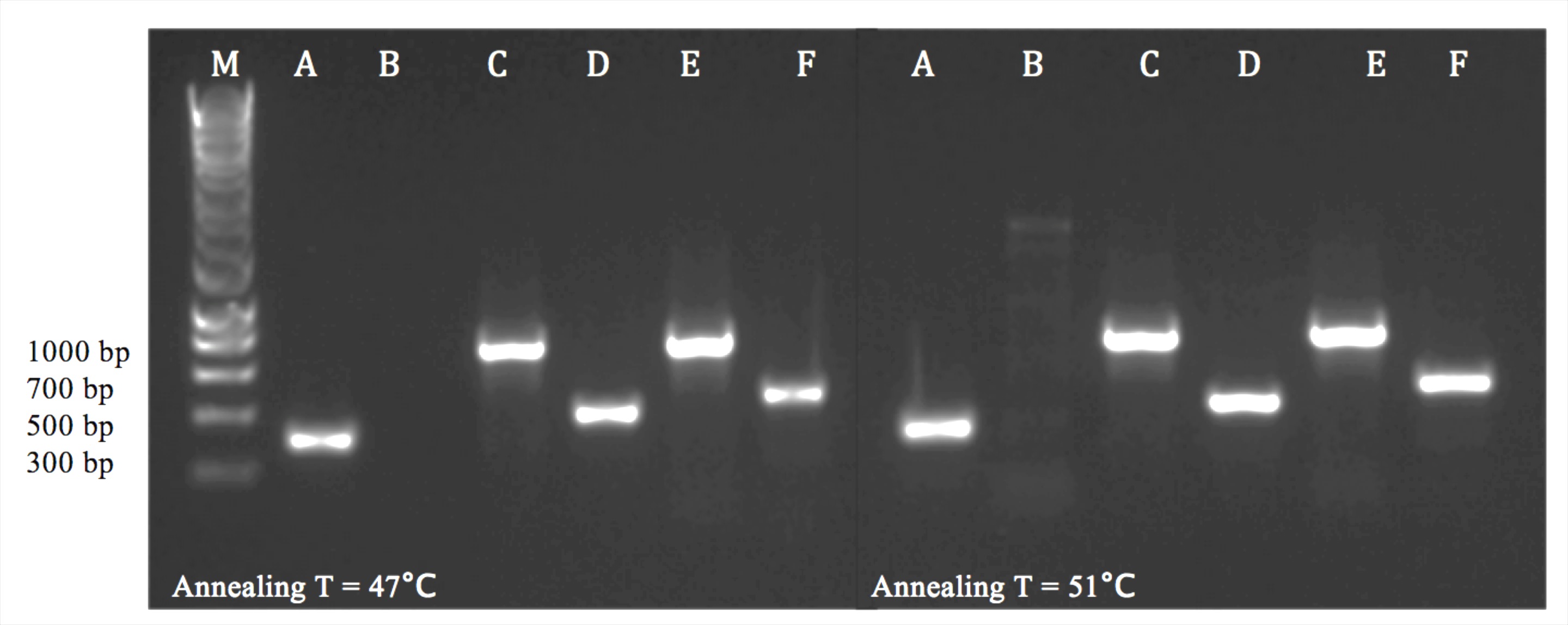

**Fig.1.** Amplification of *Chromera* ribosomal genes using newly-designed *Chromera*-specific PCR primers. M refers to the marker or DNA ladder. A refers to positive control reaction (*Symbiodinium* gDNA and *Symbiodinium*-specific primers), while B refers to negative control reaction (MQ water as a template). C, D, E and F are *Chromera* gDNA tested with the four primer pairs at two different annealing temperatures of 47°C and 51°C.

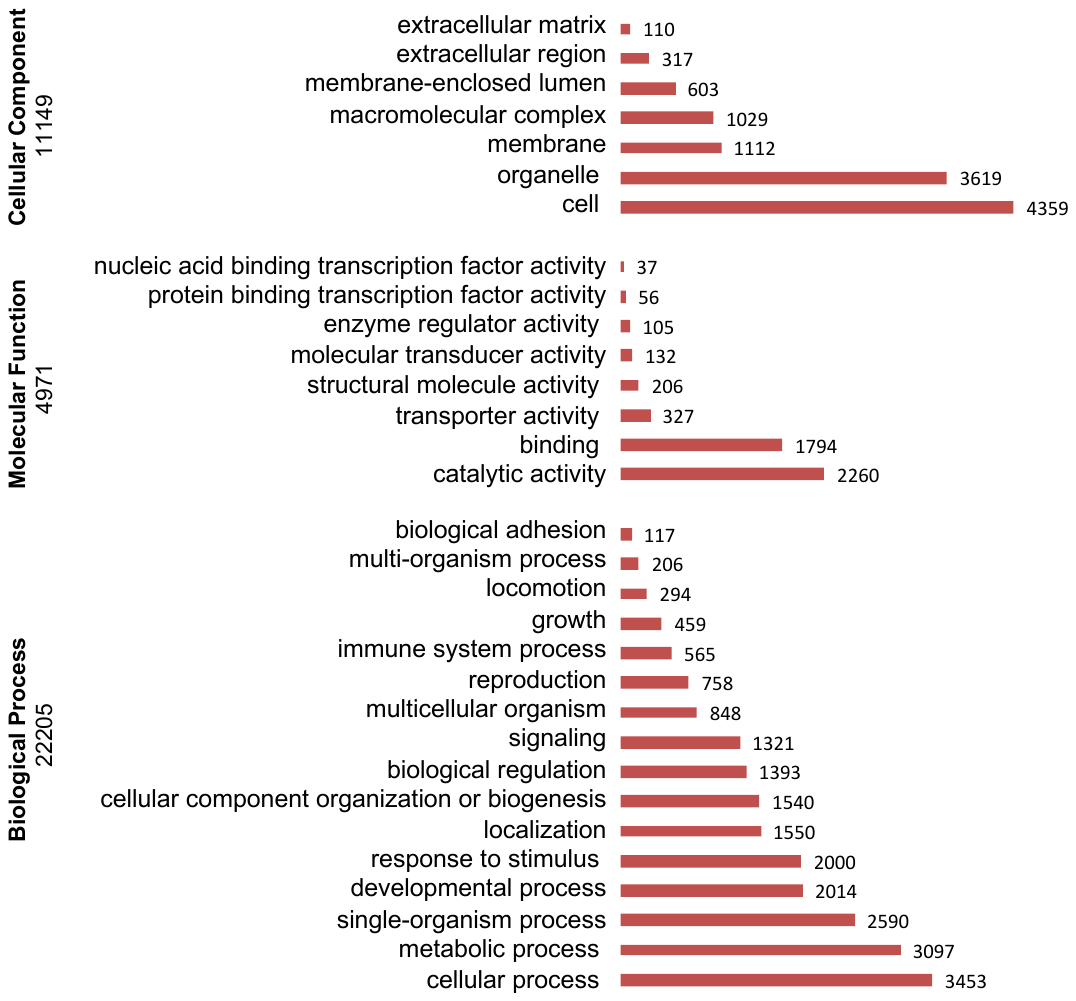

**Fig.2.** Gene Ontology (GO) assignment (2nd level GO terms) of the GBR *Chromera* transcriptome. Biological processes (A) constituted that majority of GO assignment of contigs (22,205 counts, 58.02%), followed by cellular components (C) (11,149 counts, 29.1%) and molecular function (B) (4,917 counts, 12.8%).

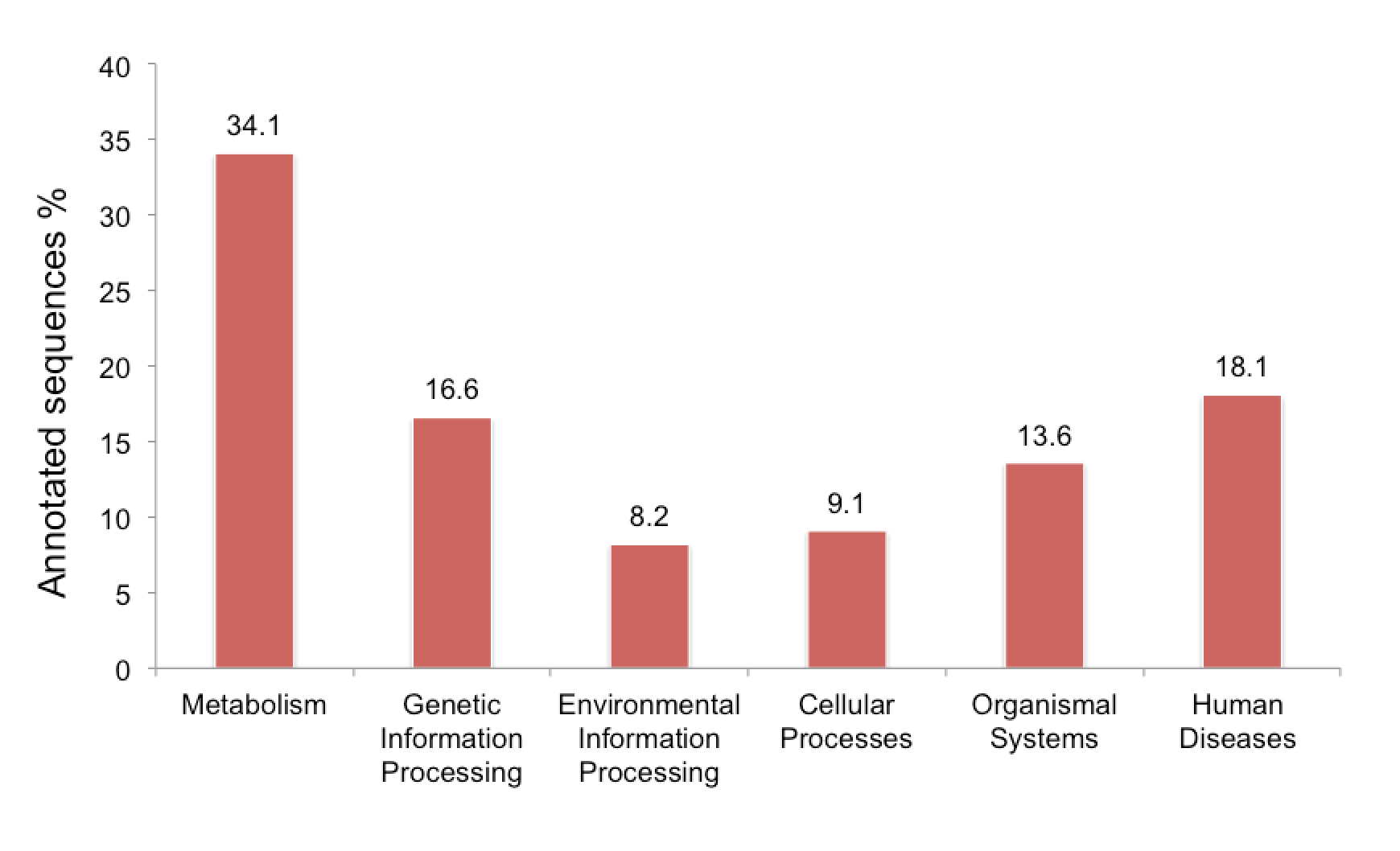

**Fig.3.** Main KEGG pathway category representation and percentages in the case of the GBR *Chromera* strain. Numbers above the bars give percent of annotated sequences in each category.

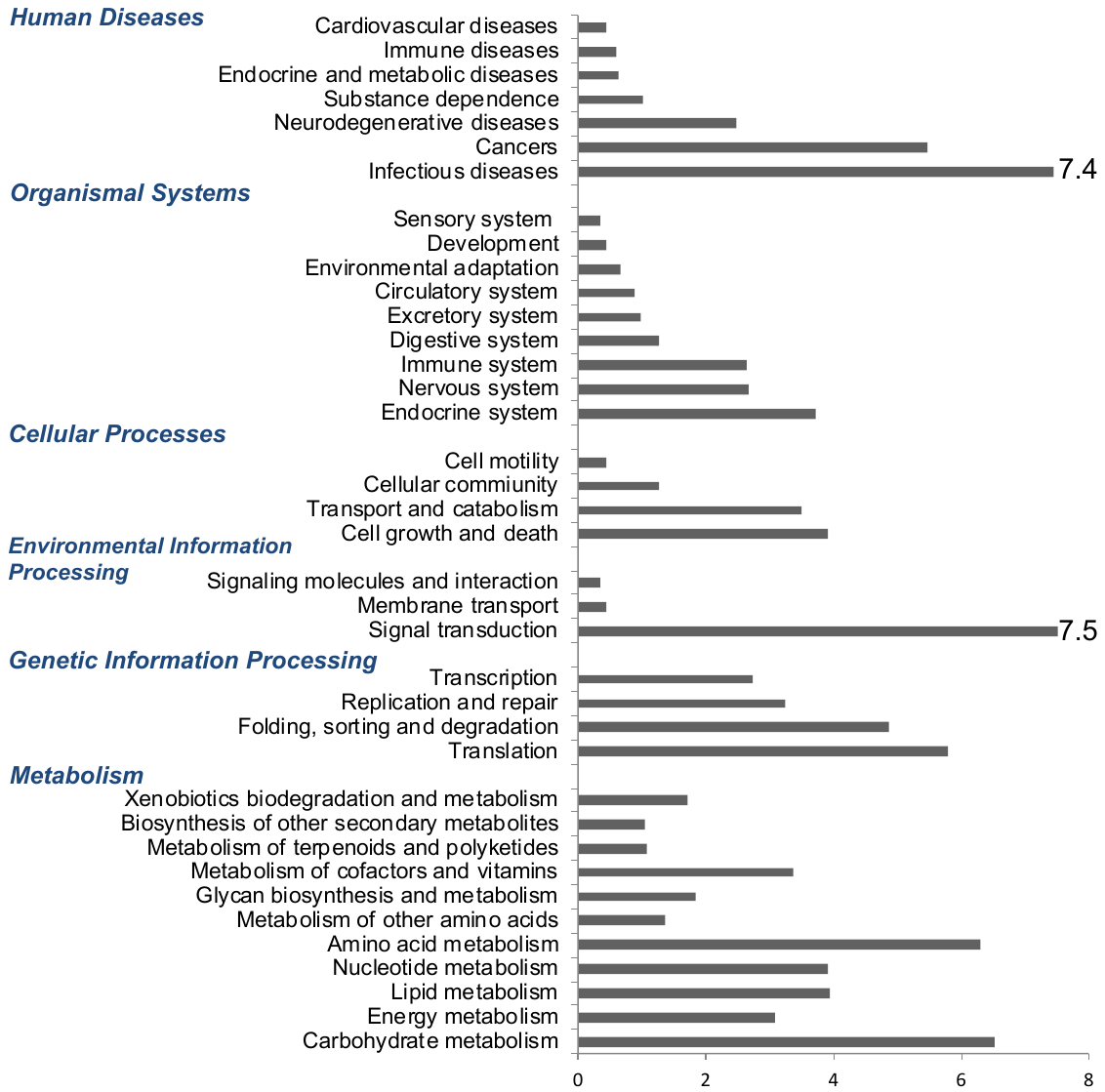

**Fig.4.** Distribution of KEGG pathways in transcriptome of the GBR *Chromera* strain. The chart shows the percentage of sequences assigned to each category.

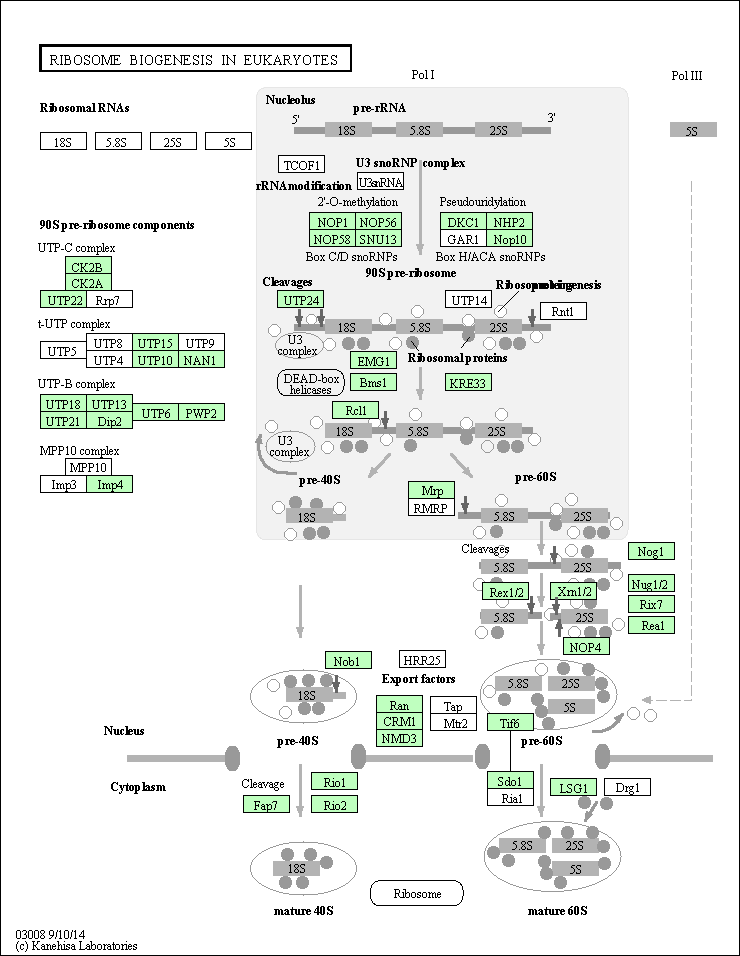

**Fig.5.** Ribosome biogenesis (eukaryotes) pathway (KO03008) identified in the *Chromera* transcriptome. KEGG pathways analysis shows *Chromera* orthologs involved in ribosome biogenesis (highlighted in green).

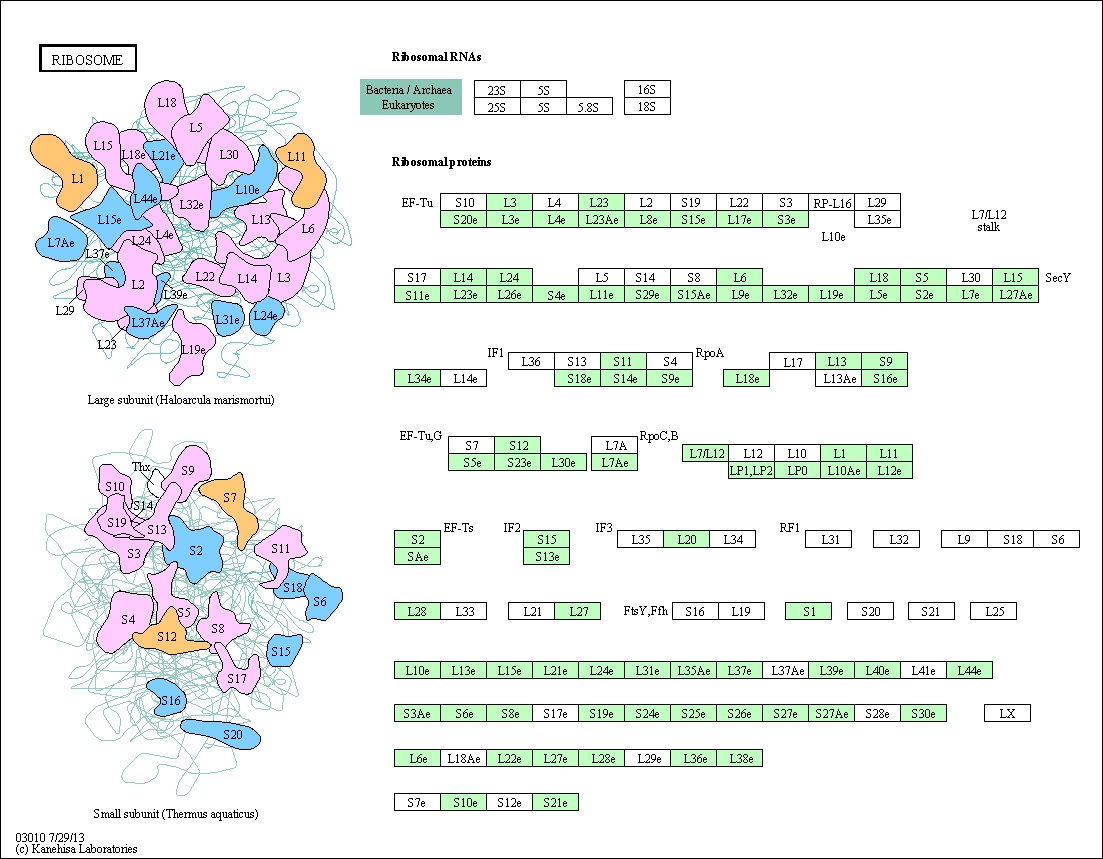

**Fig.6.** Ribosome pathway (KO03010) identified in the *Chromera* transcriptome. KEGG pathways analysis shows *Chromera* orthologs involved in ribosome (highlighted in green).

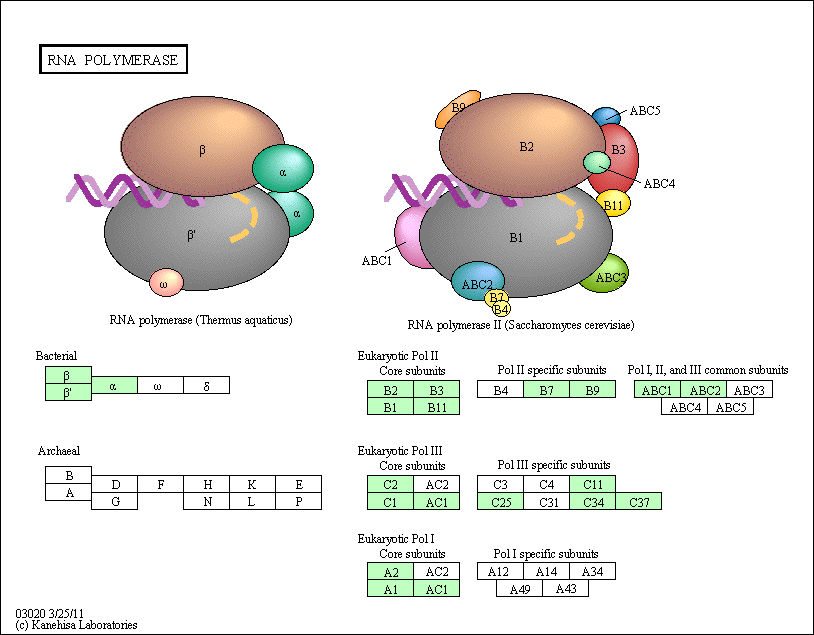

**Fig.7.** RNA polymerase pathway (KO03020) identified in the *Chromera* transcriptome. KEGG pathways analysis shows *Chromera* orthologs involved in RNA polymerase (highlighted in green).

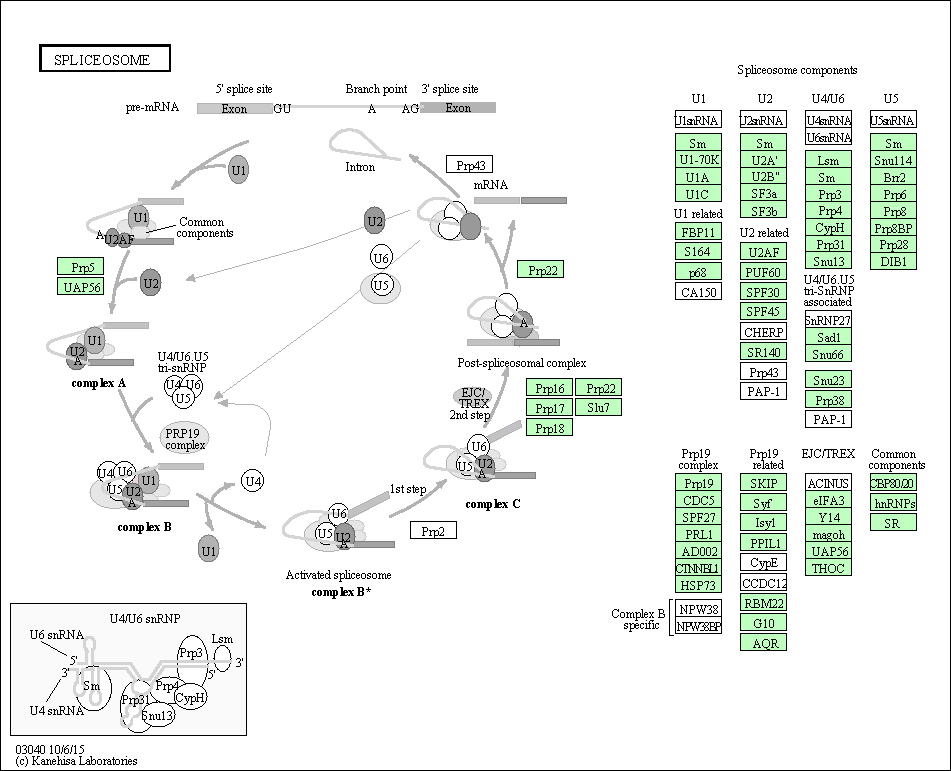

**Fig.8.** Spliceosome pathway (KO03040) identified in the *Chromera* transcriptome. KEGG pathways analysis shows *Chromera* orthologs involved in spliceosome (highlighted in green).

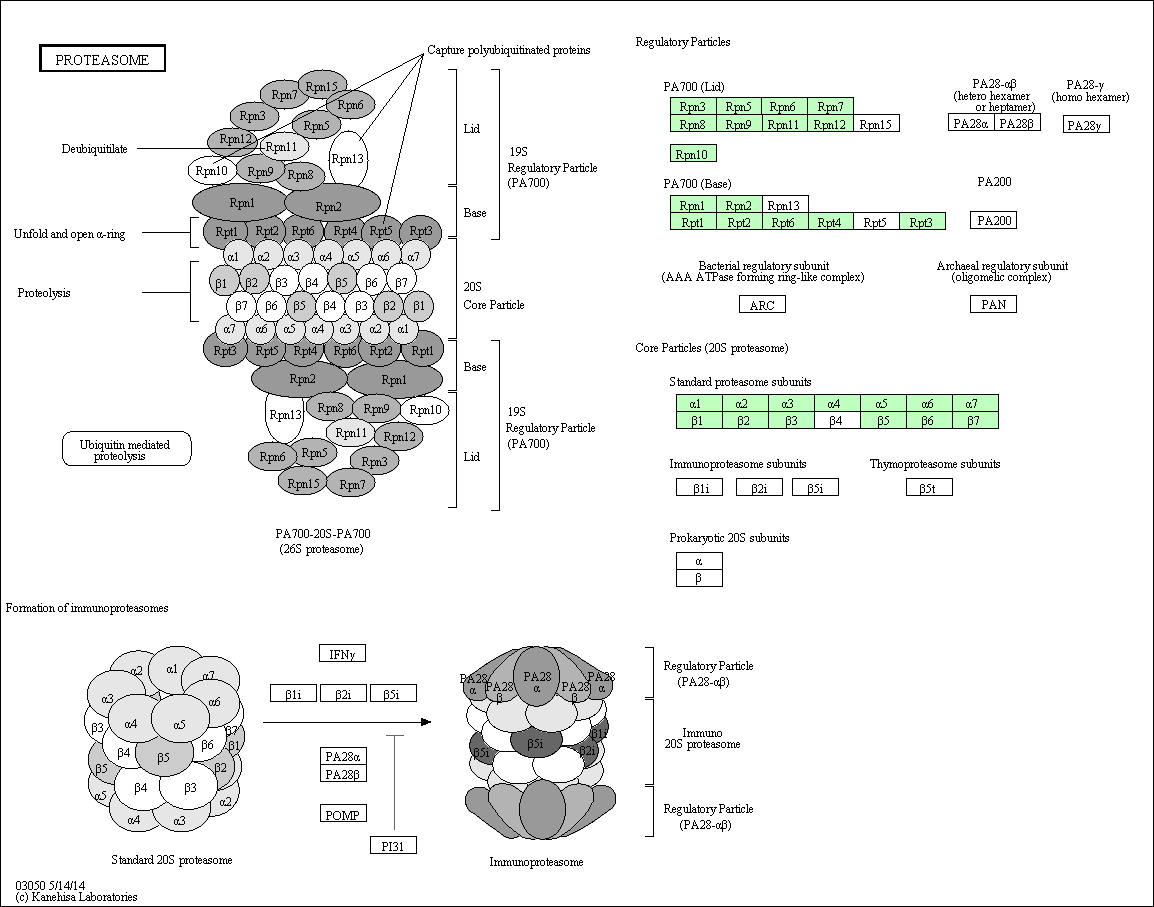

**Fig.9.** Proteasome pathway (hsa03050) identified in the *Chromera* transcriptome. KEGG pathways analysis shows *Chromera* orthologs involved in proteasome (highlighted in green).

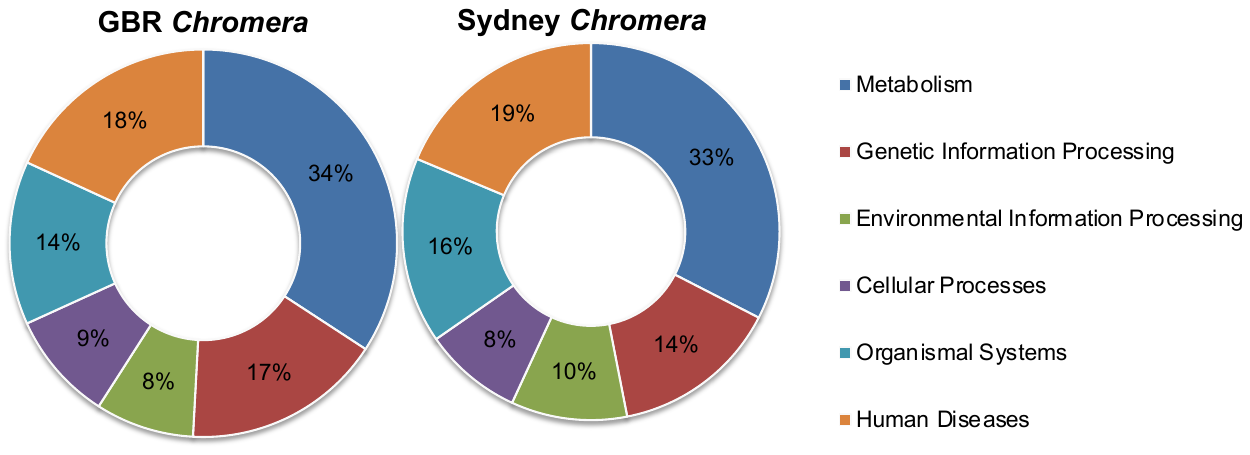

Fig.10. Overall distribution of the main KEGG categories in GBR and Sydney *Chromera*. The doughnut charts show the percentages of the sequences assigned to the six KEGG categories.

Table 1. List of *Chromera velia* specific PCR primers used for verifying the identity of the cultures. Primers were designed using the NCBI Primer-Blast tool (<http://www.ncbi.nlm.nih.gov/tools/primer-blast/>)

| Primer  Pairs | Target gene/region | Product length | Primer Sequence |
| --- | --- | --- | --- |
| 1st | LSU RNA  28S region | 755 bp | Forward Primer  **AGCCTAAGTGGGAGATCCGT**  Reverse Primer  **ACAAAGAAAGCTGCGTGCTG** |
| 2nd | LSU RNA  28S region | 416 bp | Forward Primer  **GTTTTGGAAAGCTTCGGCGT**  Reverse Primer  **ACGGATCTCCCACTTAGGCT** |
| 3rd | SSU RNA  18S region | 778 bp | Forward Primer  **CCGACTAGAGATTGGCGGTC**  Reverse Primer  **CTGACGGACTGTCGTGTGAA** |
| 4th | SSU RNA  18S region | 482 bp | Forward Primer  **TTCACACGACAGTCCGTCAG**  Reverse Primer  **CAGCACTGCAAACACATGCT** |

**Table 2.** Summary of KEGG orthology data for the GBR *Chromera* strain

| KEGG categories | No. of KO-annotated sequences (%) | No. of pathways |
| --- | --- | --- |
| Metabolism | **1442** | **127** |
| Carbohydrate metabolism | 275 (19.07) | 15 |
| Energy metabolism | 130 (9.01) | 8 |
| Lipid metabolism | 167 (11.58) | 17 |
| Nucleotide metabolism | 165 (11.44) | 2 |
| Amino acid metabolism | 266 (18.44) | 13 |
| Metabolism of other amino acids | 57 (3.95) | 6 |
| Glycan biosynthesis and metabolism | 78 (5.40) | 12 |
| Metabolism of cofactors and vitamins | 142 (9.84) | 12 |
| Metabolism of terpenoids and polyketides | 45 (3.12) | 13 |
| Biosynthesis of other secondary metabolites | 44 (3.05) | 13 |
| Xenobiotics biodegradation and metabolism | 73 (5.06) | 16 |
| Genetic Information Processing | **702** | **22** |
| Translation | 245 (35) | 5 |
| Folding, sorting and degradation | 205 (29.1) | 7 |
| Replication and repair | 137 (19.5) | 7 |
| Transcription | 115 (16.4) | 3 |
| Environmental Information Processing | **349** | **33** |
| Signal transduction | 317 (90.8) | 27 |
| Membrane transport | 18 (5.2) | 2 |
| Signaling molecules and interaction | 14 (4) | 4 |
| Cellular Processes | **386** | **20** |
| Cell growth and death | 165 (42.7) | 7 |
| Transport and catabolism | 148 (38.4) | 5 |
| Cellular community | 54 (13.9) | 5 |
| Cell motility | 19 (4.9) | 3 |
| Organismal Systems | **577** | **69** |
| Endocrine system | 157 (27.2) | 14 |
| Nervous system | 113 (19.6) | 10 |
| Immune system | 112 (19.4) | 15 |
| Digestive system | 54 (9.4) | 9 |
| Excretory system | 42 (7.3) | 5 |
| Circulatory system | 37 (6.4) | 3 |
| Environmental adaptation | 28 (4.9) | 5 |
| Development | 19 (3.3) | 3 |
| Sensory system | 15 (2.5) | 5 |
| Human Diseases | **764** | **65** |
| Infectious diseases | 314 (41) | 24 |
| Cancers | 231 (30.3) | 20 |
| Neurodegenerative diseases | 105 (13.7) | 5 |
| Substance dependence | 43 (5.7) | 5 |
| Endocrine and metabolic diseases | 27 (3.5) | 3 |
| Immune diseases | 25 (3.3) | 4 |
| Cardiovascular diseases | 19 (2.5) | 4 |

**Table 3.** Selected KEGG pathways/protein complexes identified in the GBR *Chromera* transcriptome

| Pathway/protein complex | Pathway ID | Known genes | Identified genes |
| --- | --- | --- | --- |
| Ribosome biogenesis in eukaryotes | KO03008 | 82 | 48 |
| Ribosome | KO03010 | 143 | 88 |
| RNA polymerase | KO03020 | 32 | 20 |
| Spliceosome | KO03040 | 121 | 86 |
| Proteasome | KO03050 | 48 | 29 |

**Table 4.** Recovery of BUSCO genes in *de novo* transcriptome of GBR *Chromera*

| BUSCO  (Eukaryote, n=303) |  | # genes | % |
| --- | --- | --- | --- |
|  | Single | 180 | 59.4 |
|  | Duplicated | 9 | 2.97 |
|  | Fragmented | 54 | 17.8 |
|  | Missing | 60 | 19.8 |
| BUSCO  (Alveolate-Stramenophile, n=234) |  | **# genes** | **%** |
|  | Single | 60 | 25.6 |
|  | Duplicated | 1 | 0.43 |
|  | Fragmented | 2 | 0.85 |
|  | Missing | 171 | 73.07 |
| BUSCO  (Protist, n=215) |  | **# genes** | **%** |
|  | Single | 99 | 46 |
|  | Duplicated | 1 | 0.46 |
|  | Fragmented | 2 | 0.93 |
|  | Missing | 113 | 52.55 |

**Table 5.** Genes mapped to the KEGG pathway Nitrogen metabolism “KO00910” in in GBR-*Chromera*, Sydney *Chromera*, *Cladocopium*, *Breviolum and P. falciparum.*

*Provided as excel file*

**Table 6.** Genes mapped to the KEGG pathway ABC transporters “KO02010” in GBR-*Chromera*, Sydney *Chromera*, *Cladocopium*, *Breviolum and P. falciparum.*

*Provided as excel file*
